# Cave ecosystems as drivers of viral diversity, endemism, and ecological function at a global scale

**DOI:** 10.64898/2026.11.13.766150

**Authors:** B. Barosa, A. Firrincieli, L. Gabrielli, A. Turatto, E. Lopo, D. Ghezzi, L. A. Garcia, P. E. Costantini, A. Danielli, J. De Waele, F. Sauro, M. J. Selensky, M. R. Osburn, M. Cappelletti

## Abstract

Caves are subterranean ecosystems that host diverse microbial communities involved in key biogeochemical processes. Yet, the diversity and ecological roles of viruses within these systems remain poorly understood. Here, we analyzed metagenomic data collected from 30 globally distributed cave systems, identifying 8,176 viral operational taxonomic units (≥ 10kb size) that have little representation within a recent database of environmental viruses. In addition, cave-specific environmental fingerprints are shown to be a strong driver of viral distribution, reflected in a high degree of endemism both within and across cave systems. Viral-host associations suggest that viruses are primarily associated with Pseudomonadota and Actinomycetota lineages, the most abundant groups across all samples and involved in diverse carbon, sulfur, and nitrogen metabolisms. The high proportion of lytic viruses targeting these groups suggests strong top-down viral control, with virus-mediated microbial turnover likely contributing to organic matter recycling. This is further supported by lytic viruses carrying the majority of auxiliary metabolic genes (AMGs), with functions related to enhancing viral progeny production by increasing resource availability and reallocating cellular resources toward viral replication. In this study, we highlight how cave environments shape viral ecology and unveil a still yet unknown component of the subsurface virosphere.

## 1 Introduction

Caves are widespread subterranean environments defined as voids that form and develop (speleogenesis) through diverse geomorphic processes, including dissolution, chemical weathering, melting, solidification, and mechanical erosion^1,2^. They occur within a wide range of lithologies, such as carbonates (limestone, dolostone, and marble), evaporites (gypsum and halite), volcanic effusive rocks such as basalts, Fe-rich rocks (iron ores), and silica-rich sedimentary and metamorphic lithologies (quartz sandstones and meta-quartzites)^1,3^. Although cave speleogenesis and host rock lithologies constrain to some degree the hydrogeological connectivity of these systems and the geochemical landscapes within (e.g., mineral inventories and groundwater geochemistry), caves are also influenced by the integration of other factors such as regional climate and precipitation regimes, surface land cover vegetation, surface connectivity and the presence of stygofauna^1^.

The deepest regions of cave systems are deprived of sunlight, with minimal organic inputs deriving from surface-percolating waters and streams^4^, which makes them oligotrophic ecosystems where primary production is decoupled from photosynthesis^5^. Despite these energetic constraints, caves host diverse microbial communities sustained by cooperative interactions, including syntrophic relationships that enable the complementation of metabolic capacities and the exploitation of a wide range of carbon and energy sources^6,7^. Chemolithoautotrophic microorganisms couple the oxidation of inorganic compounds, including Fe(II), Mn(II), and H₂S^8^, to CO₂ fixation, thereby supporting primary production and sustaining food webs within some cave ecosystems^9^. Trace atmospheric gases, such as H_2_, CO, and CH_4_, in aerated cave systems can also support chemolithoautotrophic growth, a process recently defined as aerotrophy^10,11^. Heterotrophic microorganisms utilize allochthonous organic matter derived from the surface as well as autochthonous organic carbon produced within the cave ecosystem^4,6,12^. Lastly, mixotrophic microorganisms are widespread, combining organic carbon assimilation with CO₂ fixation and energy acquisition from organic/inorganic compounds, reflecting an adaptive flexibility suited to the scarce and variable carbon supply of oligotrophic cave environments^11^. While these metabolic processes have been extensively analyzed and reviewed in several studies highlighting the importance of cave microorganisms (mainly bacteria) in processes with ecological and biotechnological relevance, including carbon sequestration, bioactive compound production, rock dissolution, mineral precipitation and speleothem formation^7,13,14^, the diversity of viruses and their ecological role in caves remains largely unexplored.

Viruses are the most abundant biological entities on Earth^15^ and are able to mediate the ecology and evolution of their host through cell lysis, horizontal gene transfer (HGT), and metabolic reprogramming mediated by virus-encoded auxiliary metabolic genes (AMGs)^16^. Ecosystem-wide studies of viruses and host interactions have been conducted in soils^17,18^, thawing permafrost^19,20^, deep-sea hydrothermal vents^21^, cold-seeps^22^, acid-mine drainage^23^, oil reservoirs^24^, hydraulic-fractures^25^, groundwaters^26,27^, and oceans^28^. Collectively, these studies show how viruses are important contributors to ecosystem functioning, with implications for microbial community dynamics and global biogeochemical processes. However, studies regarding cave viruses have been mostly restricted to viruses infecting cave bats^29,30^, with only one study focusing on environmental viruses in Carlsbad Caverns^31^, and limited to just 4 samples. The widespread distribution of cave systems and their substantial contribution to the subsurface biosphere, necessitate a comprehensive understanding of the microbial and viral communities they harbor at a larger-scale.

Here, we performed extensive genome- and viral-resolved metagenomics on a dataset comprising 199 metagenomes, representative of 30 cave systems worldwide, and encompassing 8 diverse speleogenetic processes. This large-scale approach allowed us to construct the first comprehensive catalogue of viral and microbial cave genomes, providing insights on the diversity and ecological roles of viruses in cave systems. Our findings expand the understanding of the Earth’s virosphere and highlight caves as major drivers of viral ecology within the subsurface.

## 2 Results

An extensive dataset from samples retrieved from diverse cave systems at a global scale (Figure 1 – A) was generated using a genome- and viral-resolved metagenomics approach (Supplementary figure 1). To minimize the influence of cave locations that may be differentially impacted by the surface environment, we manually curated the dataset to exclude samples that were either collected from regions exposed to light, or sampled within the entrance of the cave systems, keeping those from the aphotic region for downstream analysis. This curated dataset, comprising more than 1 TB of sequencing data, was obtained by coupling data newly generated in this study (52 samples) with data obtained from public databases (147 samples), including both short-reads (189 samples) and long-reads (10 samples). This resulted in a total of 199 samples from 30 cave systems (Supplementary table 1). Some caves were comprehensively sampled (e.g., Mammoth cave, USA; n= 21; Supplementary figure 2), whereas others were only represented with a single sample (e.g., Rio Garrafo cave, Italy; n= 1).

**Figure 1.**
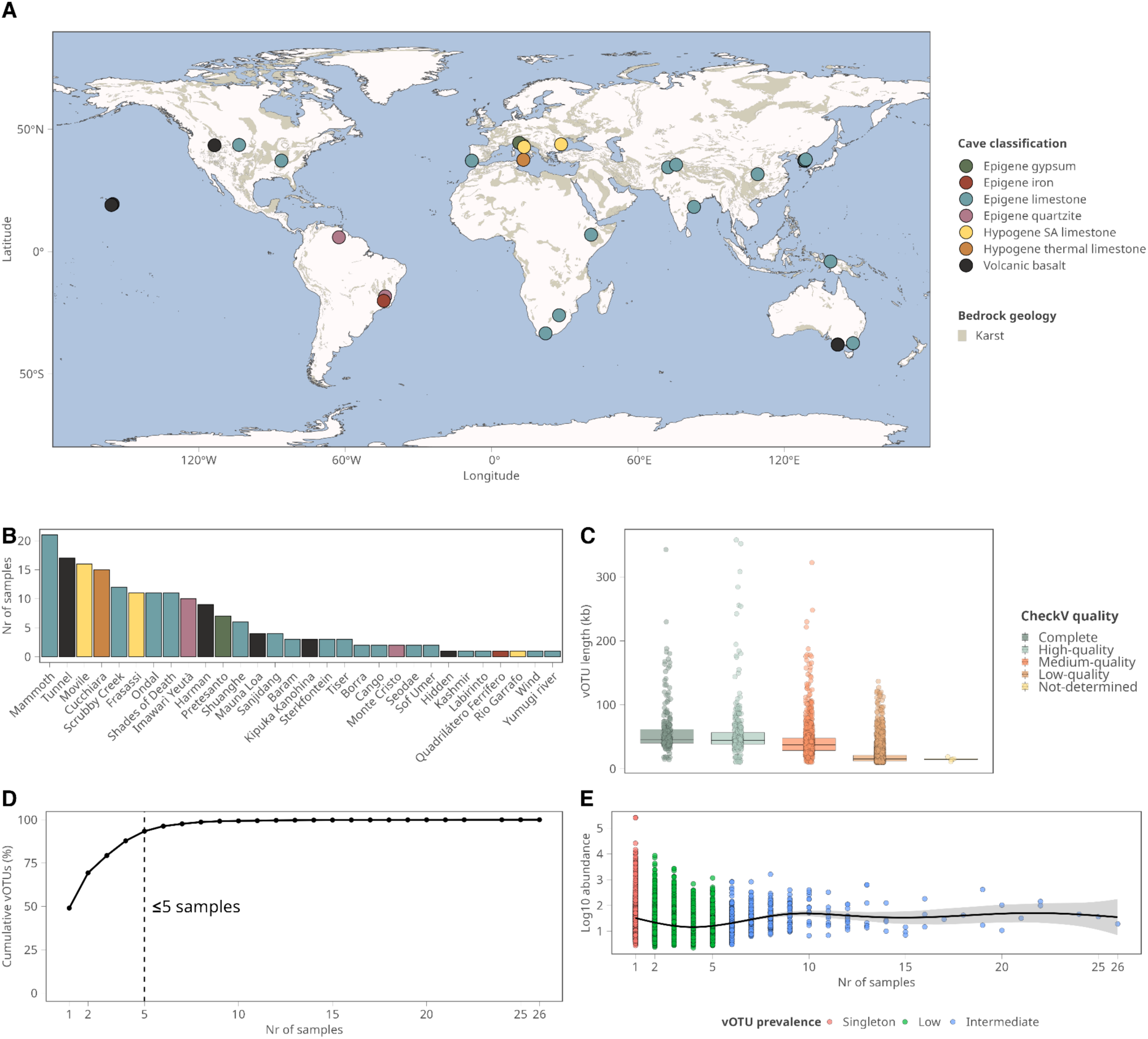
Overview of viruses in cave systems. **A)** Map illustrating the number of samples analyzed in this study representative of different cave classifications (cave classification used were adopted by the framework proposed by De Waele and Gutiérrez^1^. Detailed information of each sample is available in supplementary table 1 and 2. **B)** Number of samples per cave system for diversity analysis. **C)** vOTU length distributed over the main classification categories provided by checkV. **D)** Cumulative curve of the vOTU percentages per number of samples. **E)** Relationship between vOTU prevalence across samples and respective abundance.

We identified 15,939 putative viral contigs (≥ 10 kb) in our metagenomic assemblies using a combined approach of two viral identification tools (Virsorter2 and deepvirfinder). Given that the assemblies were from bulk metagenomes and not viral fractions (below 0.22 µm), to reduce the number of possible false positives, a set of tuning rules was applied (see methods), resulting in 10,202 viral contigs kept for downstream analysis. These were then clustered into 8,176 viral operational taxonomic units (vOTUs), representative of species thresholds (95% ANI and 85% alignment fraction). CheckV identified 275 as complete, 349 as high-quality, 1,106 as medium quality, 6,441 as low-quality and 5 as non-determined (Supplementary table 3 and Supplementary figure 3). The average size of the vOTUs in our viral catalogue is 24.48 Kb, with higher quality vOTUs having increasing lengths (Figure 1 - B). The majority of our vOTU collection was present in less than 5 samples, with 48% of vOTUs present in only one sample (Figure 1 - D), which suggests the presence of cave-specific viral diversity and a high degree of endemism, a pattern which has also been observed in other ecosystems, such as hydrothermal vents and soils^21,32,33^. In addition, there was no clear relationship between the number of samples in which a vOTU was detected and its abundance (Figure 1 - E). The accumulation curve (Supplementary figure 2) shows that our viral catalogue does not fully capture the total diversity, suggesting that additional sampling is necessary to uncover further viral diversity within cave ecosystems.

Taxonomic profiling was based on the ICTV classification and carried out using Phabox2 and geNomad. Up until the class level, 93 % of all vOTUS were assigned a respective taxonomy, however, taxonomic novelty increased at lower taxonomic ranks, with only 7.2 % of vOTUs classified at the genus level (Supplementary table 5). In addition, the majority of vOTUs were assigned to the class of Caudoviricetes within the phylum Uroviricota (n = 7787; 92 % of the total; Supplementary figure 5), which are tailed dsDNA viruses and commonly recovered from metagenomic datasets^34^. These results indicate that while a large fraction of vOTUs belong to known and established viral taxonomy, there is a high percentage of novelty in orders, families, and genera not yet represented in reference databases. In addition, the low identification rate compared to reference databases also highlights the unexplored viral diversity within cave systems.

### 2.1 Cave viruses are a novel reservoir of viral diversity

To infer the novelty of cave viruses compared to the current knowledge of the Earth’s virosphere, we leveraged the recently developed MetaVR database of environmental viruses^34^. By clustering the MetaVR data with our viral catalogue, we obtained a total of 2,953,977 species level clusters (95% ANI, with 85 % alignment fraction). Most clusters were unique to MetaVR, with 2,945,578 clusters (99.7%) containing only MetaVR vOTUs. In contrast, 7,241 clusters (0.2%) contained only cave vOTUs, while 1,158 clusters (0.04%) included vOTUs from both datasets (Figure 2 - A). The cave-associated vOTUs from these two cluster categories were subsequently used to construct a protein-sharing network with vConTACT3 to identify genus-level viral clusters (VCs; Figure 2 - A). We found a total of 5,673 VCs, from which 4,397 were singletons. Among these, 4,388 (99.7%) corresponded to cave vOTUs, indicating a high degree of novelty relative to available reference databases. Consistent with this, only one cave vOTU clustered with a NCBI Refseq reference (Supplementary table 9).

**Figure 2.**
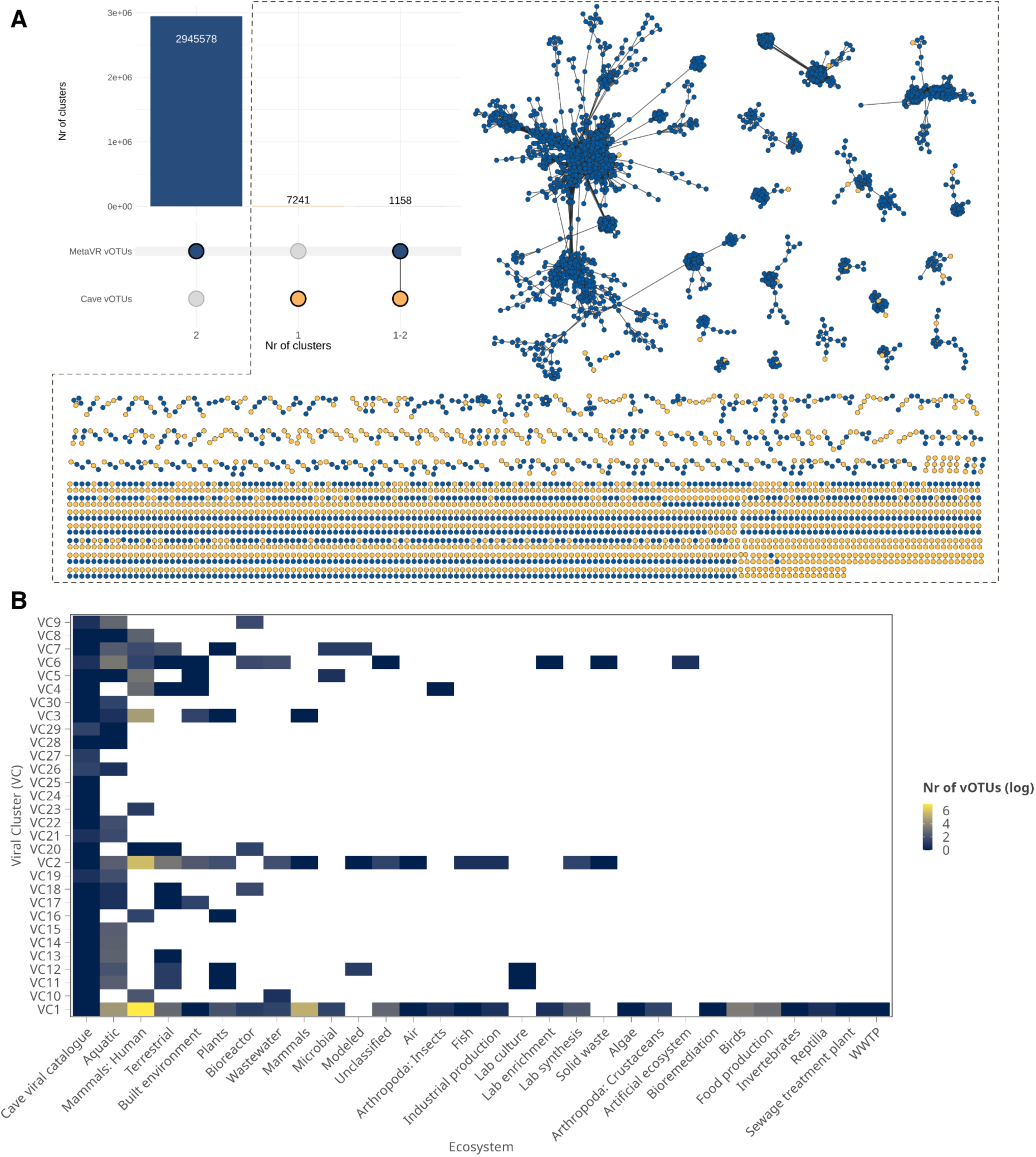
Global comparison of cave vOTUs against the MetaVR database of environmental viruses. **A)** Number of clusters of cave vOTUs against the MetaVR database, unique to MetaVR, unique to our cave viral catalogue, and clustered with shared vOTUs (singletons were removed to improve figure clarity); **B)** Biggest 20 protein clusters and the respective environments where those vOTUs were retrieved. Annotation provided within MetaVR.

Among the top 20 VCs that contained both cave and MetaVR vOTUs, the aquatic ecosystem category was the most frequent (Figure 2 - B), with 612 cave vOTUs clustering within VCs containing aquatic-associated MetaVR sequences (across all VCs). Within this broad category, we could find vOTUs retrieved from a few cave samples (Supplementary table 9), more specifically samples from a cave lake, and from other subterranean ecosystems, such as groundwaters, oil seeps, hydrothermal vents, continental margins, as well as shale gas/oil reservoirs. Surface aquatic ecosystems were also associated with cave vOTUs, including ocean, river, wetlands, intertidal zone, and creek related samples. The second most prevalent ecosystem category across the top 20 VCs was mammals-human, although only 18 cave vOTUs clustered within VCs containing vOTUs from this category. This pattern may be associated with the presence of broadly distributed human-related viruses in environmental samples in the largest VCs. Terrestrial ecosystems were the third most prevalent category across the top 20 VCs, but accounted for a substantially larger number of associated cave vOTUs (445), which suggests that terrestrial-associated viral sequences may be concentrated in lower number VCs. Within this category, the most prevalent subcategory was related to cave samples also present within our viral catalogue (wall biofilms), as well as coil crusts, tropical forest, and oil-contaminated soils. The small overlap of cave vOTUs with viruses from other ecosystems reflects a pattern of biome-specific endemism, with caves representing a vast reservoir of still unknown viruses which have a low degree of representation within current databases.

### 2.2 Ecological drivers of cave viruses

To investigate the main ecological drivers mediating viral distribution in caves, we compared viral diversity and composition across samples. Initially, to understand the broad viral biogeography we performed a distance-decay analysis (based on Bray-Curtis dissimilarities), and found significant differences in viral composition with a negative distance-decay relationship (slope = −0.013, p < 0.001; Supplementary figure 4). The regression model showed an adjusted R^2^ of 0.13, indicating that geographic distance explained approximately 13% of the observed variation. A Mantel test further indicated a significant association between geographic distance and viral community dissimilarity (Spearman’s rank correlation: p = 0.0018). The relatively low variance explained by geographic distance suggests that other factors might contribute to the viral distribution observed in cave systems. Therefore, we further explored viral diversity within the context of different cave systems and cave classification types.

We found that vOTUs assigned to Caudoviricetes were the most abundant across all cave classification types (Figure 3 - A; Supplementary figure 5 - across all samples). Conversely, the sample retrieved from an iron cave (only one sample representative) showed instead a higher abundance of giant viruses belonging to the classes Megaviricetes and Tectiviricetes. Alpha-diversity analysis based on Shannon diversity index shows that epigenic gypsum caves have the lowest mean alpha diversity values, in comparison with the hypogenic SA (sulfuric acid) limestone caves (Figure 3 - B). For dissolution derived limestone caves, hypogenic SA caves were shown to have significantly higher viral diversity compared to epigenic caves (Kruskal–Wallis rank-sum test followed by Dunn’s post hoc test, with Benjamini–Hochberg correction; adjusted p-value (p.adj) < 0.001). Lytic viruses were more abundant across all cave classifications in comparison with lysogenic ones. Additionally, we found a significant positive correlation between viral and microbial diversity (Shannon diversity; R^2^ = 0.28, p < 0.01; Figure 3 - D).

**Figure 3.**
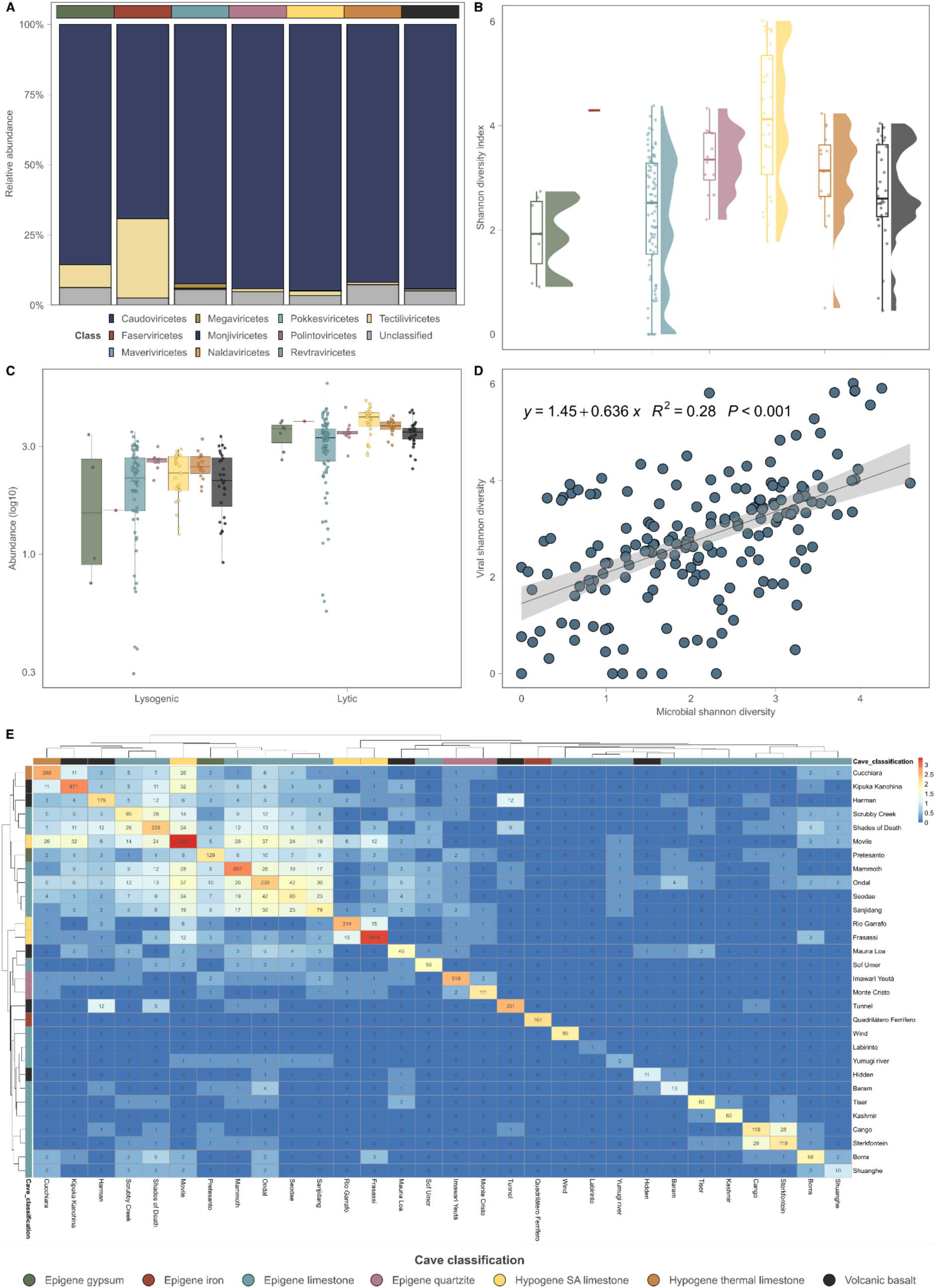
Viral taxonomy and diversity across caves. **A)** Diversity bar plot of taxonomic distribution of vOTUs across the different classification types **B)** Alfa-diversity based on the Shannon diversity index **C)** Abundance of lytic and lysogenic viruses in the different types of caves **D)** Correlation between microbial diversity (Shannon diversity index) and viral diversity (Shannon diversity index) in cave systems. **E)** Heatmap showing the number of vOTUs shared among samples within the same cave system compared with those shared across different cave systems.

To understand the broad distribution of vOTUs, we accessed their presence both within and between cave systems. We found that a substantially higher proportion of vOTUs were shared among samples belonging to the same cave system (47.9%), than among samples from different cave systems (2.9%; Figure 3 - E). Epigenic limestone caves have higher numbers of shared vOTUs suggesting that underlying speleogenesis may contribute to viral dispersion. Although cave classification had a significant effect on viral community composition, it explained only a small fraction of the observed variance (PERMANOVA, adonis2, 999 permutations, R^2^ = 0.07, p = 0.001). Individual cave systems accounted for substantially more of the variance (PERMANOVA, adonis2, 999 permutations, R^2^ = 0.27, p = 0.001), indicating that local, cave-specific fingerprints exert a stronger constraint than broader speleogenetic classification alone. In addition, beta dispersion (PERMADISP) was also significant for both variables, suggesting that part of the observed differences may be driven by heterogeneity in within-group dispersion. We observed the same pattern with the microbial communities (Cave classification, R^2^ = 0.09, p = 0.001; Cave name, R^2^ = 0.33, p = 0.001), suggesting that cave systems do also constrain the microbial distribution and diversity. Together these results indicate that viral biogeography in caves is shaped by unique cave fingerprints likely resulting from a combination of factors that also affect cave microbiota, including speleogenetic processes, surface connectivity and microclimatic conditions^35^.

### 2.3 Cave viruses primarily target hosts associated with key biogeochemical functions

To investigate viral-host associations and their potential role in microbial community dynamics and functions, we applied multiple sequence similarity-based approaches (see Methods) to screen vOTU dataset against 1,586 dereplicated (98 % ANI), medium to high quality metagenome-assembled genomes (MAGS; ≥ 50% genome completeness and < 10% contamination), recovered from the same set of samples. In total, 16.3% of vOTUs could be assigned to at least one host, a proportion similar to values reported in previous studies and indicative of the limited fraction of viral hosts that can currently be predicted across ecosystems^24,26,32,36,37^. Of the linked vOTU set, 19.03% were associated with multiple potential hosts. The predicted host MAGs spanned 38 prokaryotic phyla, including one archaeal lineage. The most abundant hosts were members of the Pseudomonadota (234 MAGs), Actinomycetota (175), Acidobacteriota (106), Planctomycetota (49), Bacteroidota (34), and Chloroflexota (Supplementary figure 6). From these, Pseudomonadota and Actinomycetota were the most abundant taxonomic lineages across all samples (Supplementary figure 6), and are also known core taxa inhabiting cave systems^6,10,38^. For this reason, we further explored the relationships between vOTUs associated with these groups.

Both the Pseudomonadota and the Actinomycetota infecting vOTUs reflect the abundance patterns of their hosts (Figure 4; Supplementary figure 7), which suggests a tight coupling between viral and microbial diversity in caves. These patterns were also observed within isolated systems such as hydrothermal vents^32^. These microbial hosts showed similar relative abundances across most cave systems at the phylum level, and their genomes encoded enzymes involved in carbon, sulfur, manganese, and nitrogen metabolisms. In addition, the overlap of some metabolic pathways across the two major lineages hints at a degree of functional redundancy, as well as the metabolic plasticity and niche breadth within these taxonomic groups. However, Pseudomonadota hosts were more associated with the sulfur cycle, as well as methanol and manganese oxidation, in comparison with Actinomycetota, which instead showed a higher number of MAGs associated with carbon fixation. In addition, we found abundant viruses associated with methanotrophs within Pseudomonadota. Caves were indicated to be greenhouse sinks, with methane oxidation activity observed within cave systems^10,39^. Thus, viral predation of methanotrophs could have a possible impact within the methane cycle in caves. Complex organic carbon degradation is also a recognized trait within these taxa, which are known to harbor functions associated with the decomposition of organic matter and with the biodegradation of complex organic molecules^40,41^. Although the surface-derived organic carbon reaching the deeper zones of caves is generally limited, sporadic and variable inputs of organic matter can access the caves depending on cave depth, surface connectivity, percolating waters^4^ and airflows, providing important energetic benefits for cave microorganisms capable of efficiently exploiting these transient resources. The predominance of lytic vOTUs associated with these hosts, together with the high abundance of Pseudomonadota and Actinomycetota across samples, suggests that these bacterial groups are subject to top-down control by viruses, with viral predation potentially contributing to the regulation of their abundance and community structure. This further entails that viral predation of these lineages can mediate and influence the turnover of abundant cave microorganisms that harbour genetic traits associated with key biogeochemical functions, influencing element cycling and organic matter recycling via the viral shunt.

**Figure 4.**
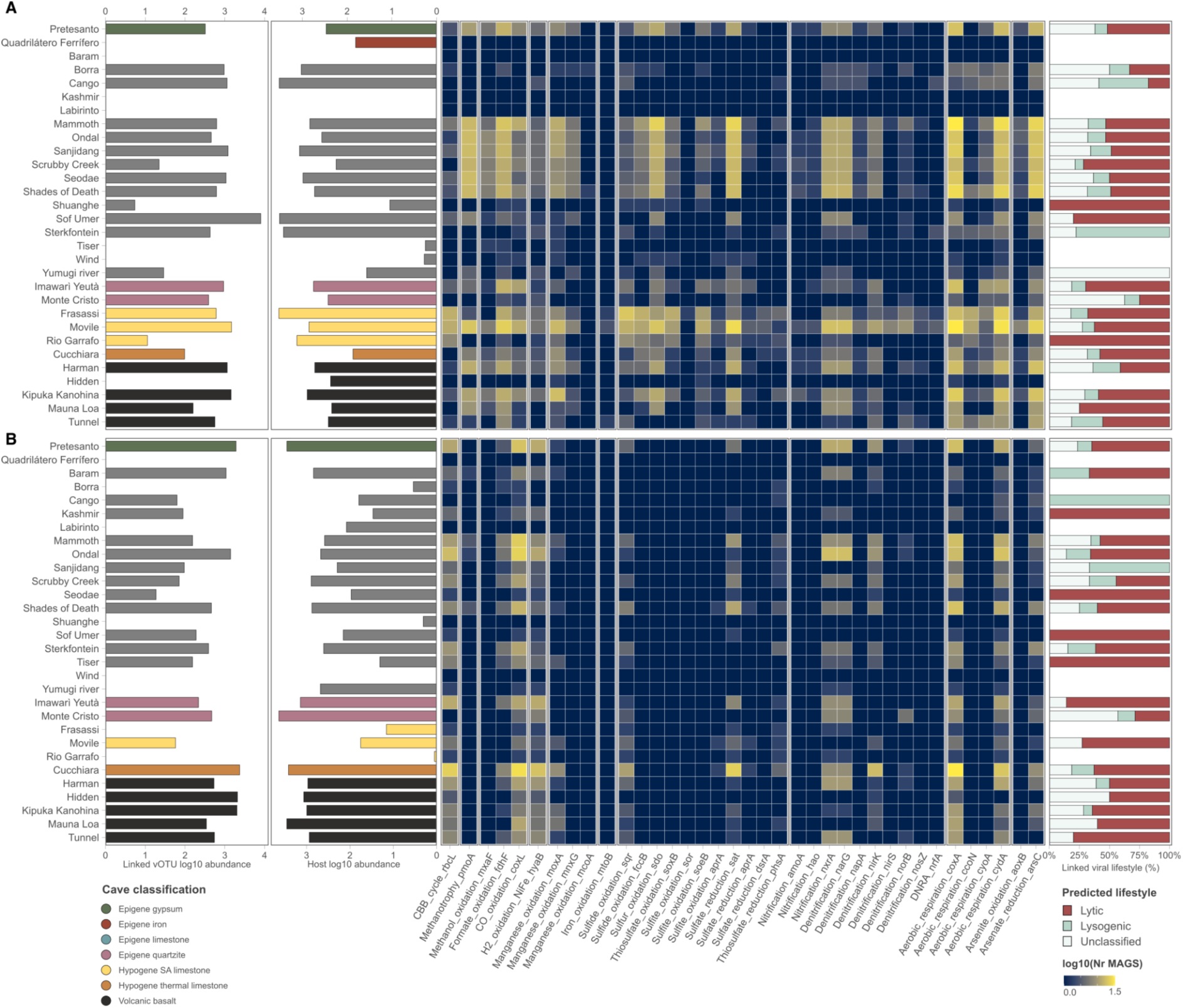
Viral interaction with the main cave bacterial phyla: **A)** Pseudomonadota and **B)** Actinomycetota. Left panels depict the viral and host abundance per cave. The middle heatmap displays the number of host MAGs carrying genetic traits associated with main metabolic pathways. Right panels refer to the respective viral lifestyles of the vOTUs identified per cave.

### 2.4 Lytic viruses in caves as the primary carriers of AMGs

To understand the role of AMGs in augmenting host metabolism we used DRAM-v, only keeping high confidence AMGs for downstream analysis (AMG confidence <=3 by DRAM-v; Figure 5 - A). We found that 302 vOTUs (3.7 %) carry high confidence AMGs, with 1,870 having been identified in previous studies, 552 experimentally validated, and 498 annotated as putative (Supplementary table 8). The major DRAM-v annotated categories were assigned to Other (n = 2247), MISC (n = 392), Organic Nitrogen (n = 178), Carbon utilization (n = 91), Energy (n = 8), and Transporters (n = 4) (Figure 5 - A). From the AMG carrying vOTUs (Figure 5 - B), the majority were present in lytic viruses (n=1,168; 17% of the total lytic vOTU set), with just very few lysogenic viruses carrying AMGs (n=16; 6.9% of total lysogenic vOTU set). To understand the distribution of these AMG-carrying lytic viruses we assessed their abundance across the different cave systems (Figure 5 - C). We found a vOTU coding for cytochrome bd ubiquinol oxidase subunit I, which has been previously observed in a hydrothermal system virus^32^. Interestingly, this virus was particularly abundant in the samples from Cucchiara cave (Sicily, Italy), a thermal limestone cave, with a cave atmosphere composed of variable concentrations of CO, CO_2_, and H_2_^42^. The presence of an AMG coding for a cytochrome with high-affinity for oxygen could provide the host with more efficient respiration and protection to various stress conditions^43^. We found widespread lytic viruses carrying AMGs related to carbohydrate-active enzymes (CAZy) across cave systems, and associated with the glycoside hydrolase families GH16 (n = 9), GH5 (n = 8), GH55 (n = 3), GH13 (n = 1), GH71 (n = 1). Overall, these GH groups are involved in breaking down complex (backbone cleavage) and simple carbohydrates (oligosaccharides) of hemicellulose, cellulose, starch, and sulfo-polysaccharides. Even though these viruses could not be linked to a specific host, the prevalent presence of CAZy within lytic viruses suggests that they promote the degradation of complex carbon substrates by their hosts, increasing energy availability and ultimately benefiting viral progeny production. The high abundance of AMGs related to amino acid metabolism, peptidases, and information systems across caves also support the rewiring of microbial cell metabolism to funnel the production of the building blocks of viral progeny. In addition, we found AMGs related to signalling, more specifically the *phoH* gene (n=59), which encodes a phosphate regulon protein induced under phosphorus starvation^23,26^. Interestingly, viruses carrying this AMG were particularly prevalent in hypogenic SA caves (present in all 3, out of 5 where they are observed). In one of these caves (e.g., Frasassi, Italy), a previous work showed that microbial biofilms are phosphorus limited and lack common phospholipids, relying instead on phosphorus-free glycolipids and betaine lipids for membrane formation^44^. Viruses carrying these genes could provide the host functions related with phosphate transport and scavenging^45^. On the other hand, we found very few AMGs associated with lysogenic viruses and restricted only to a limited number of cave systems (supplementary figure 9), and with functions associated with amino-acid and CAZy as major categories.

**Figure 5.**
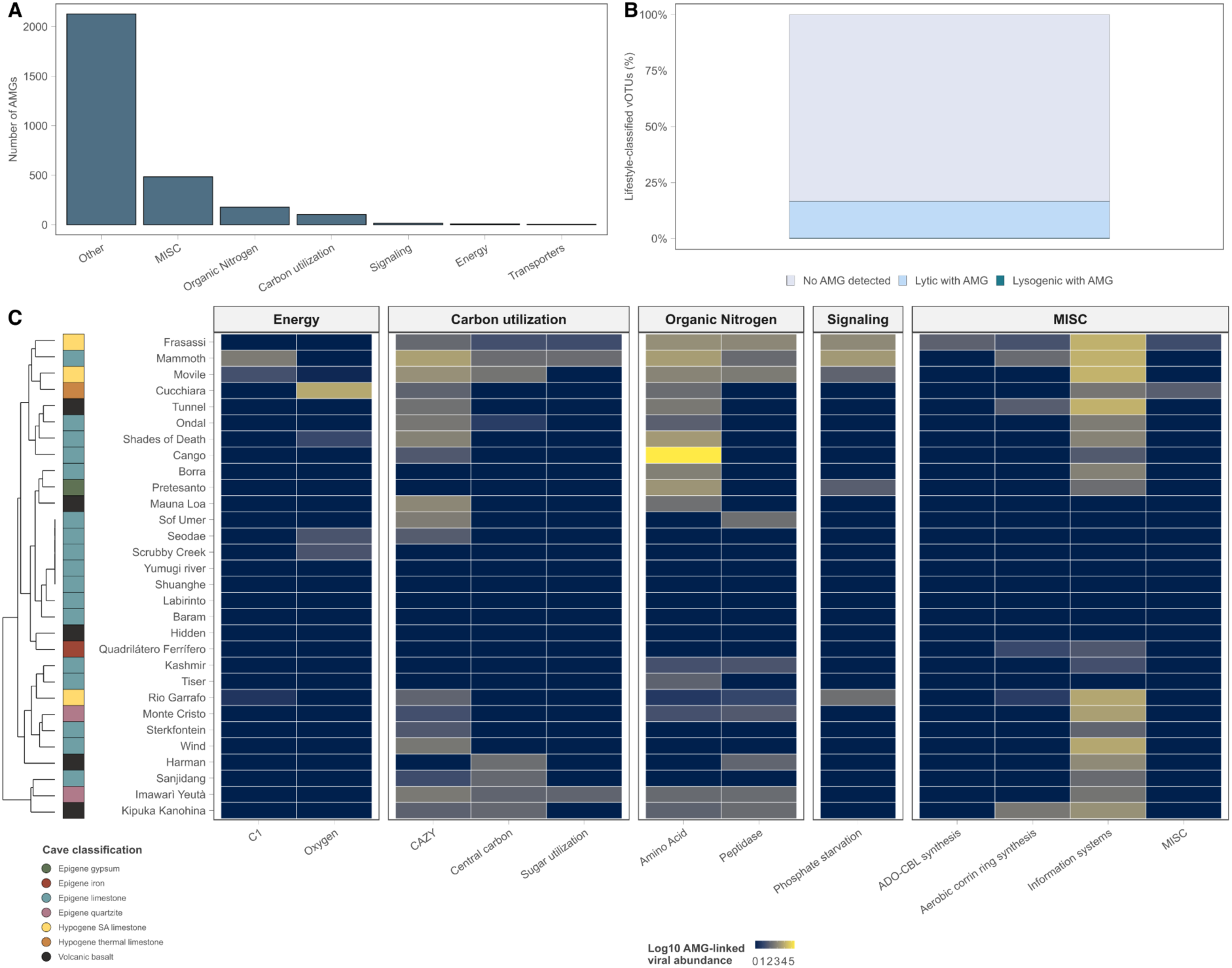
Auxiliary metabolic genes (AMGs) in the cave vOTUs. **A)** Number of total AMGs per functional category (via DRAM-v) **B)** Percentage of AMG-carrying vOTU per lifestyle **C)** Heatmap highlighting the abundance of the AMG-carrying vOTU per cave, and their respective functional annotation (via DRAM-v).

## 3 Discussion

Microbial communities are increasingly recognized as key contributors to cave ecosystem functioning, by mediating a wide range of processes, including carbon and nutrient cycling, trophic interactions, rock dissolution, and mineral precipitation^9,10,14,46–50^. By comparison, viruses remain an understudied component of cave microbiomes, with only one study investigating viral diversity and ecological roles in these systems^31^. Here, we conducted the first global-scale investigation of viral diversity across 30 caves representative of 8 different speleogenetic processes and host-rock lithologies, providing a broad perspective on the diversity, distribution, and potential ecological roles of viruses in cave ecosystems.

Most identified viruses in caves belong to the class Caudoviricetes (formerly Caudovirales) whereas classification at lower taxonomic ranks remained largely unresolved. These tailed dsDNA phages are among the most commonly detected DNA viruses across various freshwater, marine, and terrestrial ecosystems but encompass extensive genomic diversity, with closely related lineages having distinct distributions across ecological niches^51^. Despite this, comparison between cave viruses with the most recent database of environmental viruses (MetaVR^34^), revealed a large number of unknown vOTUs distinct from those detected in other environments, indicating that caves are a vast and still largely unexplored reservoir of viral diversity. This pattern is often observed in similar ecosystem-wide studies, including subsurface ecosystems such as oil reservoirs and groundwater wells and boreholes^24,26,27,52^, and have been associated with a high degree of habitat-specific viral diversity (also referred to as viral endemism). Despite this strong habitat-specific structuring, we also detected some cave viruses in surface ecosystems, suggesting that cave viral communities are not completely isolated and that some degree of connectivity may occur across surface-subsurface environments.

In contrast to some surface and subsurface ecosystems (e.g., soils^53^, acid mine drainage sites^54^, and oil reservoirs^24^), geographic distance, although significant, explains only a small proportion of the variance in viral community composition across caves. Conversely, cave identity and associated microbial community composition explained substantially larger proportions of the variation (R² = 0.33 and 0.28, respectively). Similar associations between viral and microbial community composition have been observed in other ecosystems, where they have been attributed to, at least in part, the dependence of viral distributions on host availability and diversity^24,51,54^. In caves, microbial communities are themselves shaped by local environmental conditions. Thus, viral community structuring may arise partly through environmental selection of their hosts, although direct effects of local physicochemical conditions on viruses may also contribute. Although caves share common features, such as the absence of light, resource limitation, and relatively stable microclimates, individual systems differ in geology, geomorphology, surface connectivity, and local environmental conditions (Figure 6)^1^. Together with geomorphological barriers that restrict dispersal, this environmental heterogeneity can make caves function as ecological islands, promoting distinct local communities of microbial hosts and viruses^55^.

**Figure 6.**
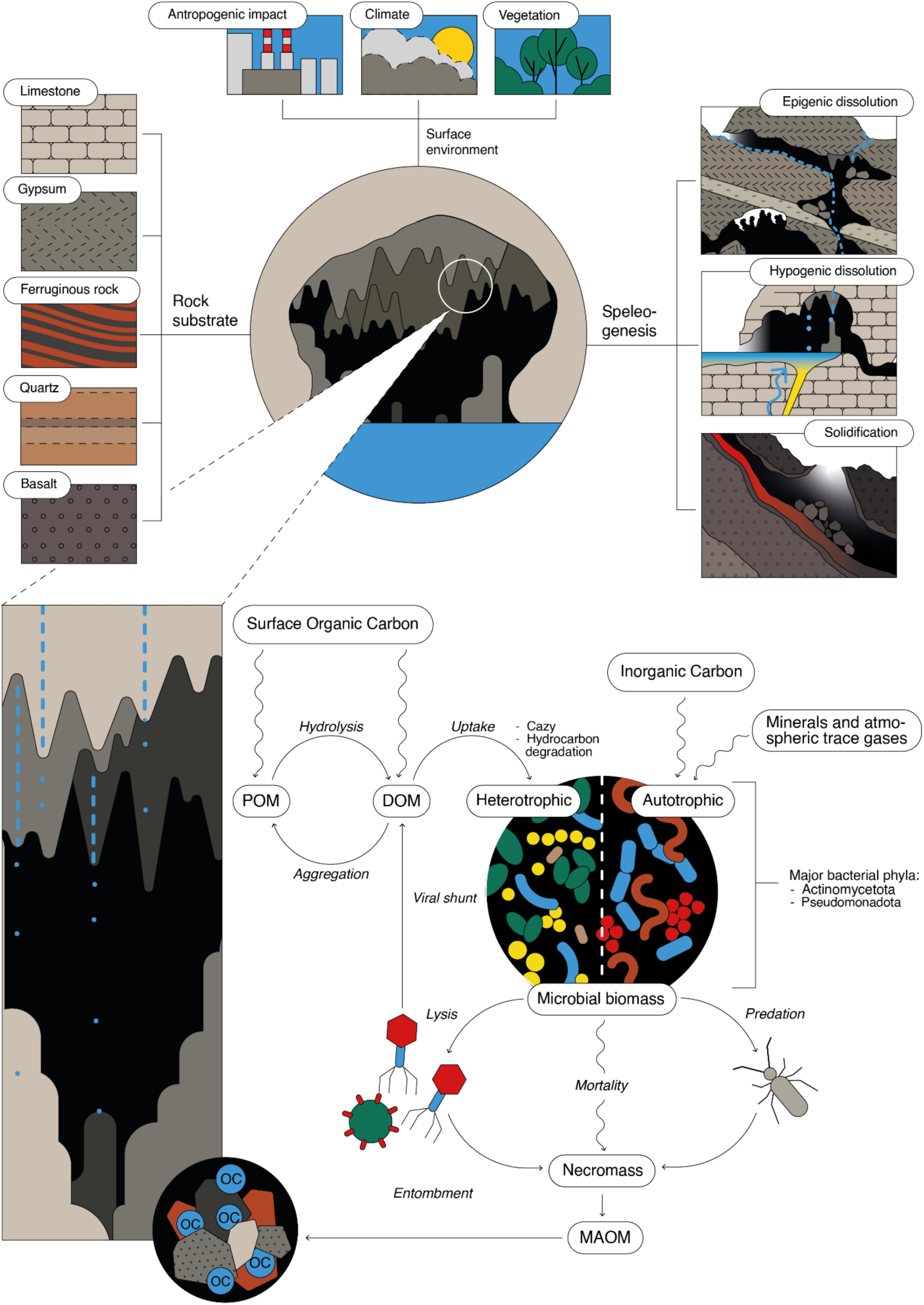
Caves as drivers of viral ecology and their associated ecological roles. The above panels refer to the main factors shaping cave unique fingerprints, whereas below is highlighted the ecological roles of cave viruses. MAOM = Mineral-associated organic matter, DOM = Dissolved organic matter, POM = Particulate organic matter. The microbial predator depicted is a Collembola, which are abundant cave arthropods, known to feed on bacteria^122^.

Consistently, vOTUs were more frequently shared among samples from the same cave than among samples from different cave systems, indicating a degree of cave-specific viral diversity. Greater connectivity within individual caves, combined with geological isolation between them, may favour within-cave dispersal while restricting viral exchange across cave systems, even when they are geographically close. This interpretation is also consistent with the viral lifestyle predictions carried out in this study, which showed a predominance of predicted lytic over lysogenic viruses across caves. Lytic lifestyles have been proposed to limit viral dispersal since free viral particles need to encounter suitable hosts, whereas lysogenic viruses can disperse together with their hosts^56^. This framework has recently been used to interpret patterns in hydrothermal vent ecosystems, where viruses found to be predominantly lytic, associated with narrow predicted host ranges, and showed largely localized distributions^32^. We found similar patterns where predicted lytic viruses predominated and ∼80% of vOTUs with assigned MAG hosts were associated with a single host. In hydrothermal vents, the most abundant viruses were associated with dominant microbial taxa, while viruses infecting low-abundance hosts were comparatively rare. In caves, the most abundant viruses were associated with the dominant bacterial groups across all samples and cave systems (i.e. Pseudomonadota and Actinomycetota). These are very broad bacterial phyla with metabolically diverse lineages involved in biogeochemical cycling and mineral transformations^8^. In particular, we show that the host MAGS belonging to these groups code for a high diversity of metabolic pathways involved in carbon, sulfur, and nitrogen metabolisms. In addition, we identified enzymes related to the oxidation of trace atmosphere gases, a process recently defined as aerotrophy^10^ across the host genomes, which included gene functions associated with methane and hydrogen oxidation. Given that most viruses predicted to infect these abundant and metabolically important groups were assigned a lytic lifestyle, they may exert top-down control on cave microbial communities through the selective infection and lysis of dominant hosts. By actively controlling the most abundant microbial populations, cave viruses could contribute to maintaining microbial diversity by allowing the establishment of less abundant and slow growing taxonomic groups, following the kill-the-winner hypothesis^57^. Thus, virus-host interactions in caves could influence not only microbial community composition and diversity but also the metabolic potential underlying biogeochemical cycling and atmospheric gas consumption and fluxes.

Viral-induced cell lysis may also promote viral shunt, releasing cellular organic matter (OM) and nutrients that can be recycled by other microorganisms. Under the oligotrophic conditions typical of cave environments, this process could provide an important source of substrates for microbial growth. Moreover, the OM and microbial necromass resulting from viral lysis that is not rapidly consumed by microorganisms can interact with reactive mineral surfaces, promoting the formation of mineral-associated organic matter (MAOM). Due to its association with minerals and occlusion within aggregates, this form of organic carbon is relatively resistant to microbial degradation. On the one hand, MAOM could be hypothesized to contribute to indirect biological mechanisms of speleothem formation by providing nucleation surfaces for mineral precipitation^58^. On the other hand, this association with minerals may favour carbon stabilization, as described in soils, where MAOM contributes to long-term carbon storage^59^, raising the possibility of a similar, yet largely unexplored, carbon reservoir in subterranean environments.

We also found that lytic viruses were the primary carriers of auxiliary metabolic genes (AMGs) in caves, suggesting their potential to reprogram host metabolism to support viral replication and progeny production. For instance, by carrying a gene encoding subunit I of the cytochrome bd terminal oxidase, cave viruses may provide functions that support host respiration under variable O₂ concentrations and enhance protection against different environmental stresses^43^. In addition, viruses carrying CAZy-related genes were particularly abundant across multiple cave systems. These genes may modify host carbohydrate metabolism during infection, potentially influencing the turnover of complex organic carbon derived from sporadic surface inputs^19,59,60^. The widespread occurrence of AMGs associated with amino acid metabolism, peptidase activity, and information-processing functions further suggests that cave viruses may reprogram host cellular processes to support the production of energy and molecular building blocks required for viral genome replication and progeny formation^61^. Finally, the presence of AMG-*phoH* genes, particularly in sulfidic caves, suggests a link between the low phosphate availability characteristic of some of these systems^44^ and a potential role in enhancing host responses to phosphate limitation. *phoH* has been reported as an AMG in viruses from diverse ecosystems^62,63^, although a direct relationship between environmental phosphate availability and the occurrence of phoH-carrying viruses remains unclear.

In conclusion, our study provides the first comprehensive assessment of viruses in cave ecosystems, highlighting their largely unknown diversity and possible role as drivers of microbial communities in subterranean environments on a global scale. Despite these insights, several methodological limitations must be acknowledged. The usage of DNA metagenomes excludes RNA viruses, which could also play key roles within cave systems as is the case of marine ecosystems^36,64^. In addition, our analyses are based on genomic inference, and for that reason they do not reflect functional activity. The overall small fraction of lysogenic viruses, host predictions, and AMGs may partly reflect genome incompleteness and the limitations of current databases and prediction tools^65^. Finally, our study relies on bulk metagenomes rather than virome fractions, which could potentially improve viral recovery^66,67^, although contrasting results have been reported^67,68^. Nevertheless, our results are consistent with one of the few previous cave virome studies, conducted across water pools in Carlsbad Caverns^31^, which similarly revealed diverse and novel viral communities dominated by Caudoviricetes and lytic lifestyles, with AMGs primarily associated with viral production. Despite the different approaches, the consistency of these findings suggests that both bulk metagenomes and virome fractions can recover comparable patterns of cave viral diversity, and that processes previously observed within a single cave system may reflect broader ecological patterns across cave ecosystems.

## 4 Materials and Methods

### 4.1 Sampling, DNA extraction and sequencing

For the samples collected at the Imawarì Yeutà (Venezuela), Cucchiara (Sicily, Italy), and Prete Santo caves (Emilia-Romagna, Italy), DNA was extracted using DNeasy PowerLyzer PowerSoil Kit, according to the manufacturer’s instructions. For the samples collected from Mammoth cave (USA), DNA was extracted using the ZymoBIOMICS DNA MiniPrep kit, according to the manufacturer’s instructions, and sequenced at NU Seq Core Facility (Northwestern University, Illinois, USA) for shotgun metagenomic sequencing on a NovaSeq 6000 SP flow cell (Illumina). For other cave systems analyzed here, raw metagenomic data were downloaded from NCBI sequence read archive (SRA) database^69^ (sample descriptions, SRA and Bioproject accessions available in supplementary table 2)^10,70–80^. The cave classification framework used here is available in detail within the supplementary information.

### 4.2 Processing of metagenomic sequence data

The raw sequencing data used in the present study was obtained using both short-read and long-read sequencing technologies (Illumina and Nanopore, respectively). For the Illumina, short-reads were quality filtered and trimmed using bbtools (https://jgi.doe.gov/data-and-tools/bbtools/), and assembled using MetaSPAdes (v.4.2.0)^81^ with default parameters. The assembly output was further investigated using metaQuast^82^ and read-mapping carried out using BBmap (v.39.33)^83^. For Nanopore, long-reads were trimmed with chopper (v.0.12.0)^84^, assembled using metaFlye (v.2.9.6)^85^, and read-mapped with minimap2 (v.2.3.0)^86^. Metagenomic binning on the individual assemblies were performed using a combination of high-performance binners^87^: Semibin2 (v.2.2.0)^88^, Metabat2 (v.2.18)^89^, and Metadecoder (v.1.2.1)^90^. The consensus refined MAGS from the three tools were obtained using MAGScoT (v.1.1)^91^, and further dereplicated using dRep (v.3.6.2)^92^ with default settings (obtaining metagenomic operational taxonomic units (mOTUs) at 98% ANI level). Completeness and contamination estimates were carried out using CheckM2 (v.1.1.0)^93^, with only medium to high-quality MAGS retained for downstream analysis (completeness > 50% and contamination < 10%).

### 4.3. MAG taxonomic and functional predictions

Taxonomic classification of the MAGs was conducted using the genome taxonomy database with GTDB-tk (v.2.5.1)^94^. Protein coding genes (ORFs) were identified using Prodigal (v.2.6.3)^95^. The predicted genes were further annotated using Kofamscan^96^ and filtered to retain KOs related marker genes involved in key biogeochemical functions (https://github.com/BernardoBarosa/biogeo_metabolic_markers). Hits with E-values ≤ 1 × 10^−20^ and bit scores above the KEGG HMM-specific thresholds were retained. When multiple hits were available for the same gene, the best hit was selected based on the lowest E-value and highest bitscore. Genes that did not pass this initial filtering were considered for rescue if they had E-values ≤ 1 × 10^−10^ and scores ≥ 80% of the KO-specific HMM threshold, following the bit-score relaxation heuristic logic implemented in Anvi’o (anvi-run-kegg-kofams)^97^.

### 4.4 Viral contig identification

Viral contigs were identified in the metagenomic assemblies using a combined approach coupling two tools: VirSorter2 (v2.1; parameters: --min-length 10000 bp, --keep-original-seq, --min-score 0.5)^98^ and DeepVirFinder (v1.0; parameters: score ≥ 0.9 and p < 0.05)^99^. The quality of the putative viral contigs was assessed using CheckV (v1.0.3)^100^. To improve the confidence of the identified viral contigs, a set of tuning rules^101^ was applied on the output of CheckV by removing viral contigs where: (i) the number of viral genes equals zero and the number of host genes ≥ 1; (ii) the number of viral genes is 3x lower than the number of host genes and provirus field is annotated as FALSE; (iii) the number of host genes is > 50 and provirus field is annotated as FALSE; (iv) contig length is > 50 kb and the number of hallmark genes is ≤ 1. The viral contigs that passed these rules were then clustered into viral operational taxonomic units (vOTUs), using the scripts provided within checkV (95% average nucleotide identity (ANI) and 85% alignment fraction). To uncover the uniqueness of our viral catalogue, we initially clustered our vOTUs with the vOTUs (≥ 10kb) included within the recently released MetaVR database^34^ using vclust^102^. To optimize resource usage, the pre-filtering step was carried out using the flags --min-kmers 20 --min-ident 0.95 --batch-size 100000 --max-seqs 10000 --kmers-fraction 0.2. The clustering step was carried out at 95% average nucleotide identity (ANI) and 80% alignment fraction. The percentage of clustered vOTUs was calculated with reference to post-filtered vOTUs, and the clustered groups were then used to compute a protein similarity network within vcontact3^103^. For visualization purposes only, the network file (*.cyjs*) generated via vcontact3 was reduced in size by eliminating clusters not including cave vOTU, and non-informative intra- and inter-genus edges. Edge removal was performed with the minimum spanning tree function *mst* from the igraph R package (v2.2.0). For intra-genus edges, a minimum spanning tree was computed guaranteeing cluster connectivity with the minimum number of edges (n − 1 per cluster), while additionally retaining all intra-genus edges with a distance ≤ 0.15 to preserve visual cohesion within clusters. For inter-genus edges, only those with a distance ≤ 0.10 were retained, in order to display strong relationships between clusters while removing weak, non-informative connections.

### 4.5 Estimating abundances across-samples

The abundance of vOTUs and their associated microbial hosts is a key metric for characterizing the diversity and distribution of viral communities in ecosystem-wide studies. This step becomes particularly relevant when datasets integrate multiple sample types and sequencing platforms, which can introduce various biases, including compositional effects and differences in sequencing depth. Even though no single abundance metric is universally optimal, a previous study has reported that coverage-based metrics provide improved performance over read count-based approaches^104^. Taking these considerations into account, in this study we used trimmed mean coverage values calculated by coverM (v. 0.7.0; 5% - end trimming to account for spurious high coverage values, with 50 % contig coverage)^105^, following community-established standards (90% read mapping identity and 75% minimum read aligned length)^106,107^. The values were then further normalized by sequencing depth (total number of quality-filtered reads per hundreds of millions of reads). Both vOTU and MAG abundance was calculated using this method (for the MAGs, a value of 25 % contig coverage was used instead). Due to the advantages of using long-reads to recover vOTUs^108^, nanopore samples were used only for vOTU generation and were excluded for abundance calculation and diversity analysis.

### 4.6 Viral taxonomic and functional annotations

Viral taxonomic affiliations were carried out using PhaBox2 (v.2.1.3)^109^ and GeNomad (v.1.11.2)^110^. The output from both tools was considered for the final consensus classification taking into account that: i) vOTUs were assigned “unclassified” if both tools failed to assign, or in instances where there was a disagreement between both tools within and below a specific taxonomic level, and ii) vOTUs were assigned to the taxonomic level determined by one of the tools if the other showed less resolution. Putative auxiliary metabolic genes (AMGs) were identified using DRAM-v (v.1.5.0)^111^. Since DRAM-v relies on a specific output of VirSorter2, all vOTUs were first processed with VirSorter2 using the flag --prepare-for-dramv. To increase confidence in AMG assignments, only genes with an auxiliary score ≤ 3 were retained, that is, characterized by the presence of viral hallmark/viral-like genes flanking both sides of the AMG. To reduce false positives, genes annotated with a viral flag (V), transposon flag (T), viral-like peptidase flag (P), or attachment flag (A) were removed^19^. Additionally, putative AMGs lacking a gene ID or functional description were discarded. AMGs passing these thresholds were then divided into known (K flag by DRAM-v; previously identified in the literature), experimentally verified (E flag by DRAM-v; been shown in a host to provide a specific function in an experimental setting), and putative AMG (M flag by DRAM-v). Viral lifestyles (lytic and lysogenic) were identified by coupling the results of checkV^100^ and VIBRANT^112^. Mismatch between both tools was assigned as “Unclassified”.

### 4.7 Host-prediction

To predict host-virus linkages, we followed multiple in-silico approaches^17,19^. Briefly, i) the vOTU sequences were compared to the MAGs generated from the same sequencing reads using BLASTn (parameters: -dust no -perc_identity 70)^113^. vOTUs with a bit score ≥ 50, E-value ≤ 10^−3^, and ≥ 70% identity across ≥ 2,500 bp were retained for downstream analysis; ii) to identify CRISPR spacers present in our MAG collection, we used MinCED (v.0.4.2)^114^. After removal of spacer sequences containing ambiguous bases, we searched them against all vOTUs using BLASTn^113^ with the BLASTn-short option and default parameters, with subsequent filtering for matches with 100% sequence identity; iii) tRNAs genes in our vOTUs were predicted using tRNAscan-SE (v.2.0.12)^115^, using bacterial/archeal models (flags -B and -A, respectively). The identified sequences were then compared to the MAG collection using BLASTn^113^, with matches with full-length alignments containing ≤2 nucleotide differences and 100 % identity retained.

### 4.8 Statistical analyses

All statistical analysis and visualization were carried out using R software (v.4.5.2)^116^. Alpha diversity was calculated using the Shannon diversity index^117^ within the *phyloseq* package^118^, with differences assessed using Kruskal–Wallis test. Differences in viral community composition among the different cave classification and cave systems was carried out using PERMANOVA, *adonis2*, with 9999 permutations, and dispersion accessed using *permadisp* in the *vegan* package (v.2.7.2)^119^. Data wrangling was carried out with tidyverse (v.2.0.0)^120^ and plotting with ggplot2 (v.4.0.2)^121^.

## 5 Data Availability Statement

The data and code supporting the findings of this study will be made publicly available upon publication (including sequencing data - SRA accessions).

## 6 Conflict of Interest

The authors declare no conflict of interest.

## 7 Acknowledgments

The authors wish to thank CINECA for awarding the ISCRA-C grant, which provided access to the Galileo 100 high-performance computing cluster to perform the analyses. The collection and analysis of samples from Cucchiara has been conducted within the Convention for the study of cave systems in the Riserva Naturale Integrale “Monte San Calogero (Kronio)” that involves the Assessorato Regionale Beni Culturali - Soprintendenza Beni Culturali ed Ambientali di Agrigento, Dipartimento Regionale dello Sviluppo Rurale e Territoriale, Servizio per il Territorio di Agrigento, Earth and marine Science Department (DiSTeM) of the University of Palermo, Department of Pharmacy and Biotechnology (FABIT) of the University of Bologna, Ass. La Venta, and Società Alpina delle Giulie (Sez. di Trieste del Club Alpino Italiano), Comm. Grotte E. Boegan. The E.L. PhD scholarship is financed by the funds PNRR-DM118/2023.

## 8 Funding

B.B.’s postdoctoral fellowship was partly funded by the PRIN 2022 project 2022EFNWME. E.L.’s PhD scholarship was supported by PNRR-DM118/2023 funds. Sequencing analyses of samples collected from the Prete Santo gypsum cave were funded by the Federazione Speleologica Regionale dell’Emilia Romagna (FSRER). Sequencing analyses of samples collected from Imawarì Yeutà cave were partly supported by University of Bologna (UNIBO) RFO funds. Mammoth Metagenomic Sequencing was performed with a seed grant to MRO from NCKRI; MRO acknowledges funding from CIFAR Earth 4D program where she is a fellow.

## 9 Author Contribution

B.B., F.A., and M.C. conceptualized the project. D.G., M.J.S., E.L., A.T., M.R.O., and M.C. contributed to sample collection, and/or DNA extraction and sequencing. A.D. and P.E.C. supported the taxonomic analysis. M.R.O. provided information on samples and associated data from Mammoth Cave. F.S. and J.D.W. contributed to cave classification and nomenclature. B.B., L.G., and L.A.G. performed the metagenomic and viral analyses. A.F. refined the network analysis and visualization. B.B. conducted data validation, curation, analysis, and visualization. M.C. provided funding for the sequencing of samples collected in Imawarì Yeutà, Cucchiara, and Prete Santo. M.R.O. provided funding for the sequencing of samples collected in Mammoth Cave. M.C. supervised the project. B.B. and M.C. wrote the first draft of the manuscript. All authors contributed to manuscript editing, and approved the final version.

## Notes

### Competing Interest Statement

The authors have declared no competing interest.

