## Supplementary information for "Cave ecosystems as drivers of viral diversity, endemism, and ecological function at a global scale"

### Cave classification

Cave classification applied in this study relies on the framework proposed by De Waele and Gutiérrez[1], with categorization based on main speleological processes and rock lithologies. Dissolution caves develop within soluble rock substrates, such as carbonates (limestone and dolostone), evaporites (gypsum and halite), and low soluble substrates, such as silica-rich sedimentary and metamorphic lithologies (quartz sandstones and meta-quartzites) and ferruginous rocks (iron ores)[1]. Within dissolution-based speleogenesis, the formation of the cave system is driven by the chemistry and movement of percolating waters within rock pores, small fractures, conduits, and bedding planes. When meteoric waters infiltrate from the surface and percolate downwards, the resulting caves are classified as epigene, regardless of lithology. Instead, when the waters have deep origins, and ascend towards the surface, often after undergoing heating, mixing, or chemical modification, they are deemed hypogene in origin[2]. Solidification caves, on the other hand, develop during the cooling of molten rock, and are called volcanic caves. Following this classification, we grouped 30 caves within 8 different categories representative of different speleogenetic processes and main host lithologies: Epigene gypsum, Epigene limestone, Epigene Quartz, Epigene Iron, Hypogene thermal limestone, Hypogene SA (sulfuric acid) limestone, and volcanic basalt caves.

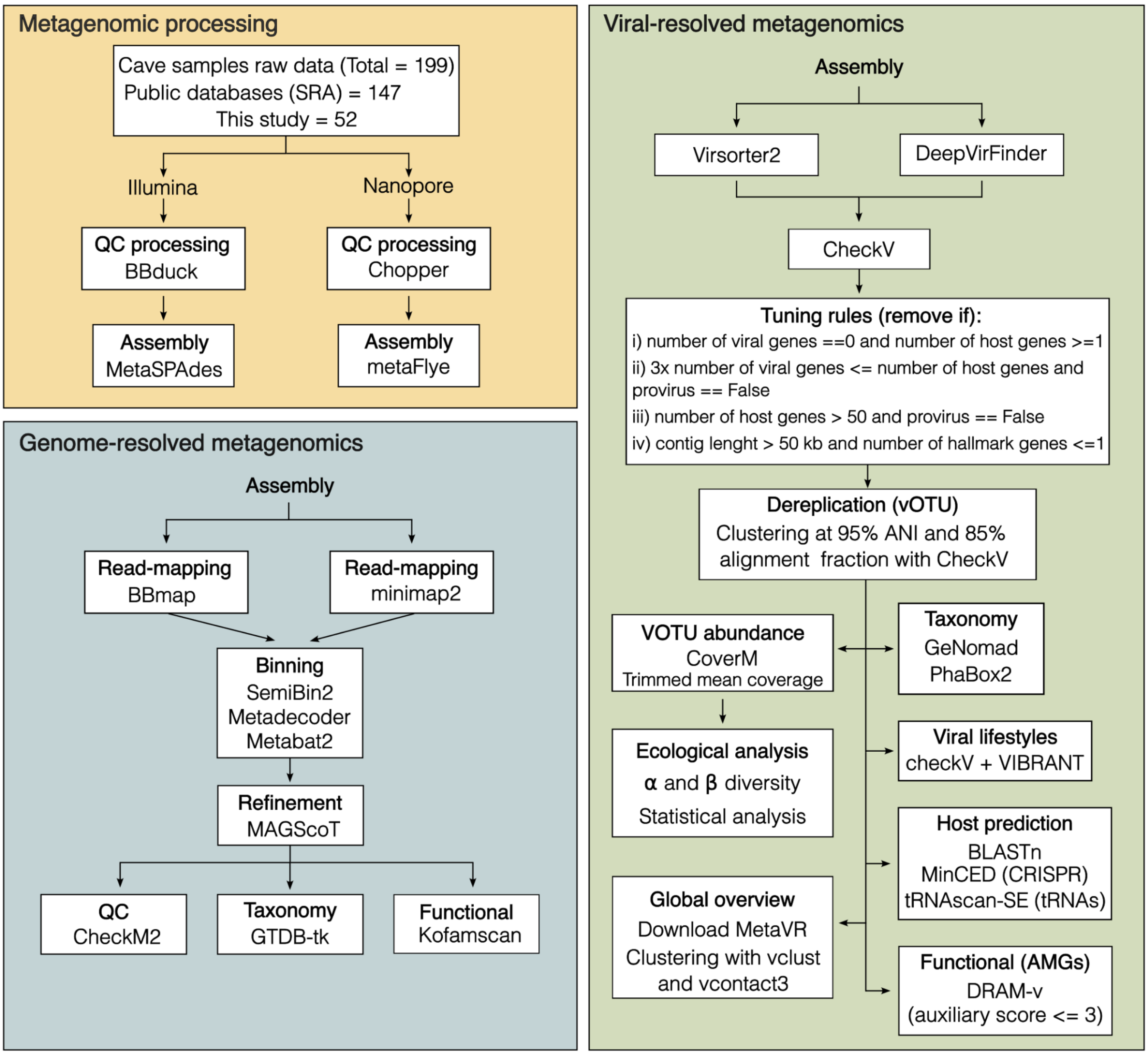


**Supplementary figure 1:** Diagram illustrating the pipeline used in the study. The three colours represent the main steps of the metagenomic processing, genome-resolved and viral-resolved metagenomics.

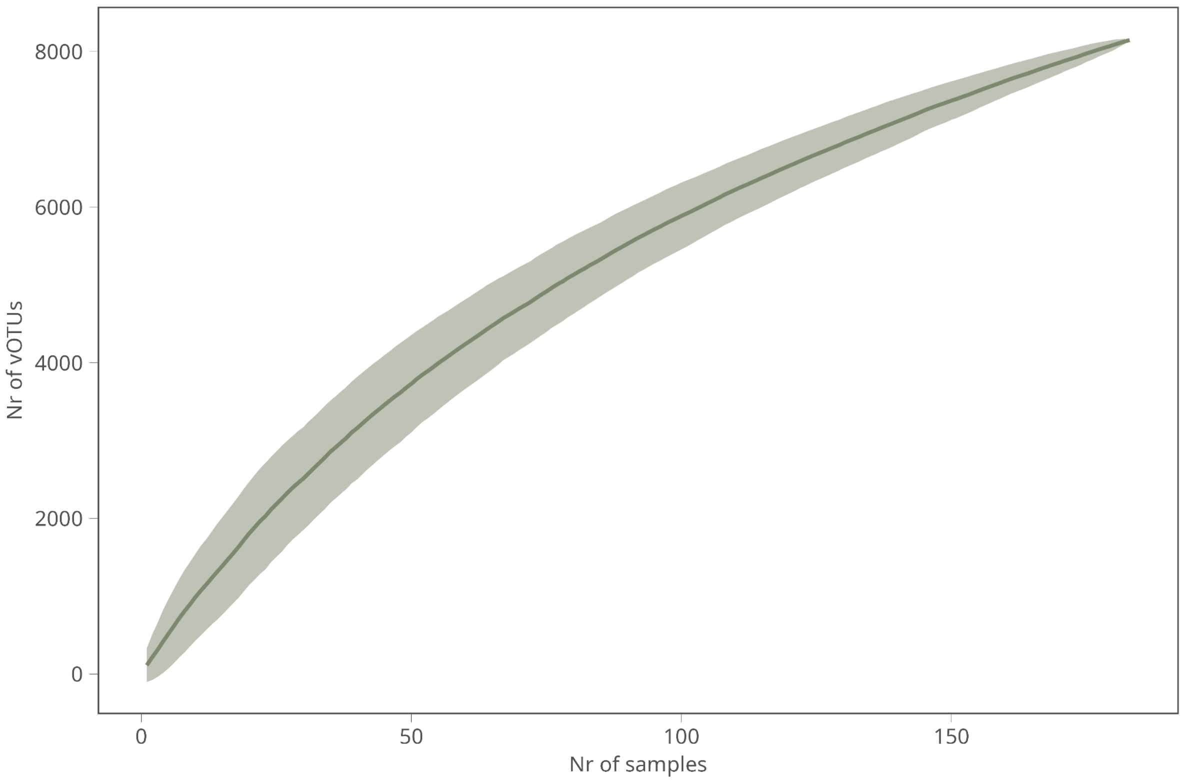


**Supplementary figure 2:** Accumulation curve of the identified vOTU as the sampling effort increased. The error bar range is represented by the grey-coloured thickness of the curve.

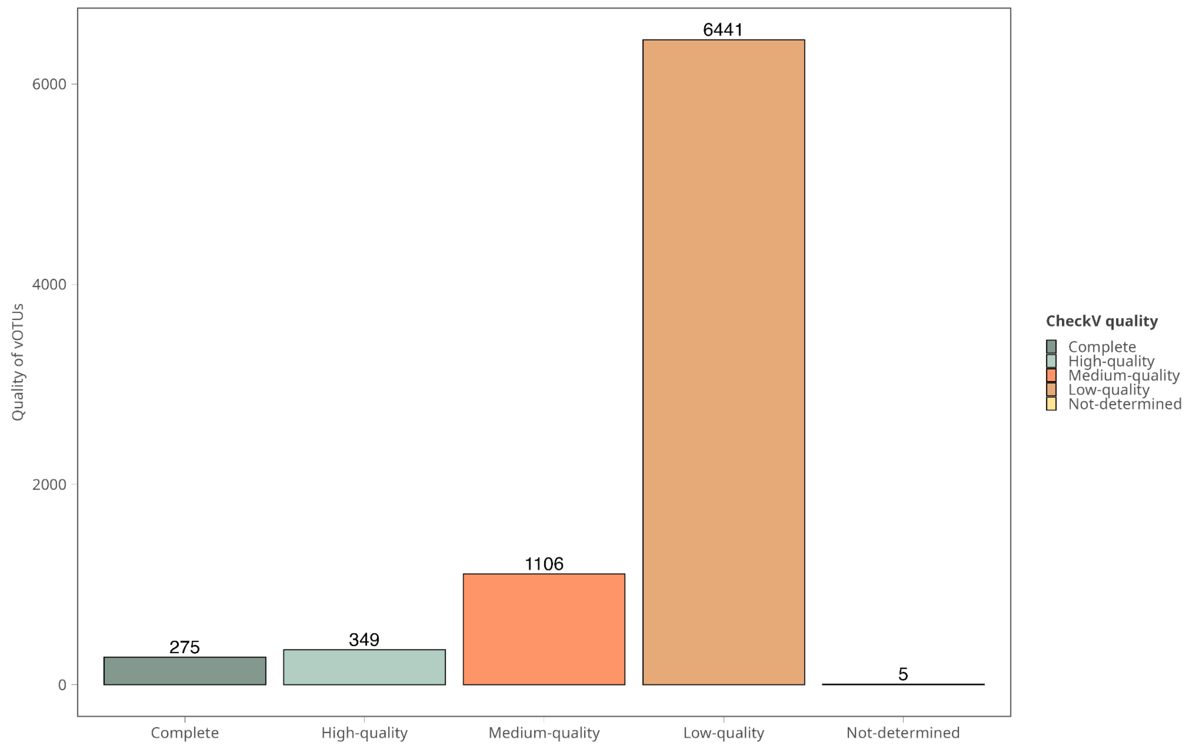


**Supplementary figure 3:** Number of vOTUs per CheckV quality metrics

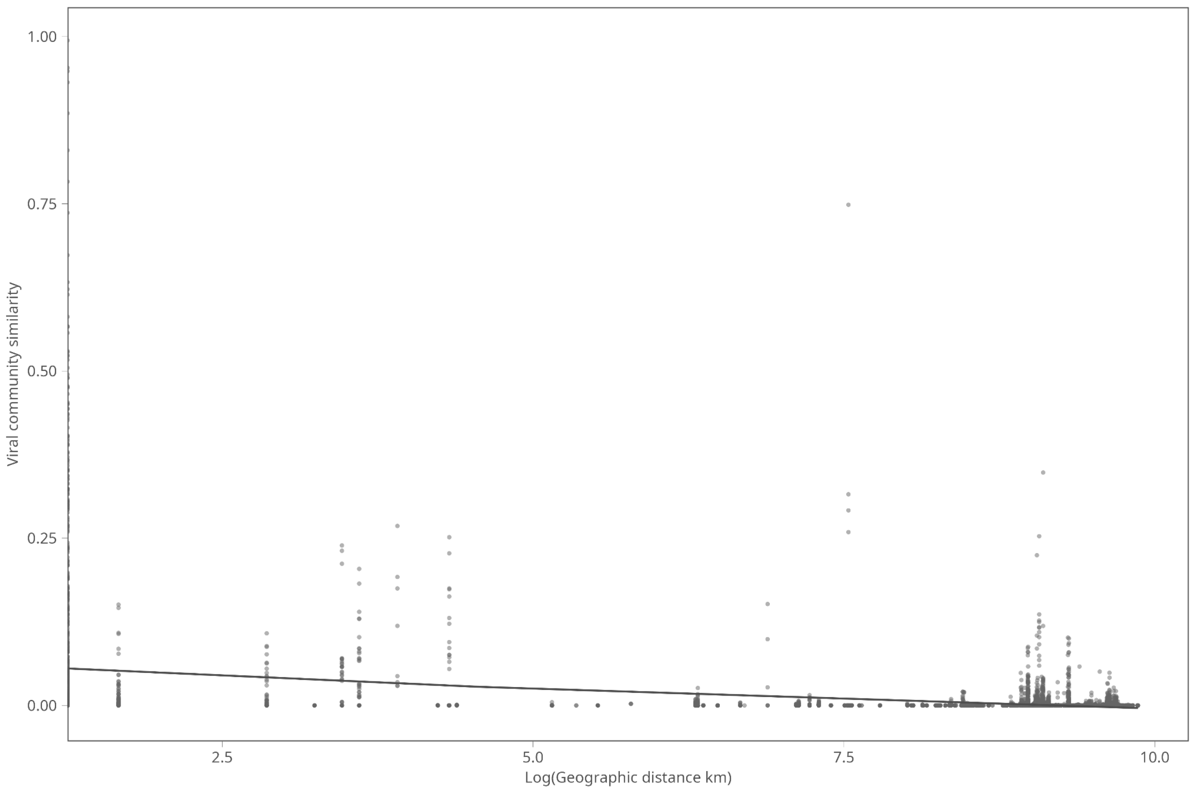


**Supplementary figure 4.** Distance-decay relationships based on Bray-Curtis similarity with linear-regression fitting line.

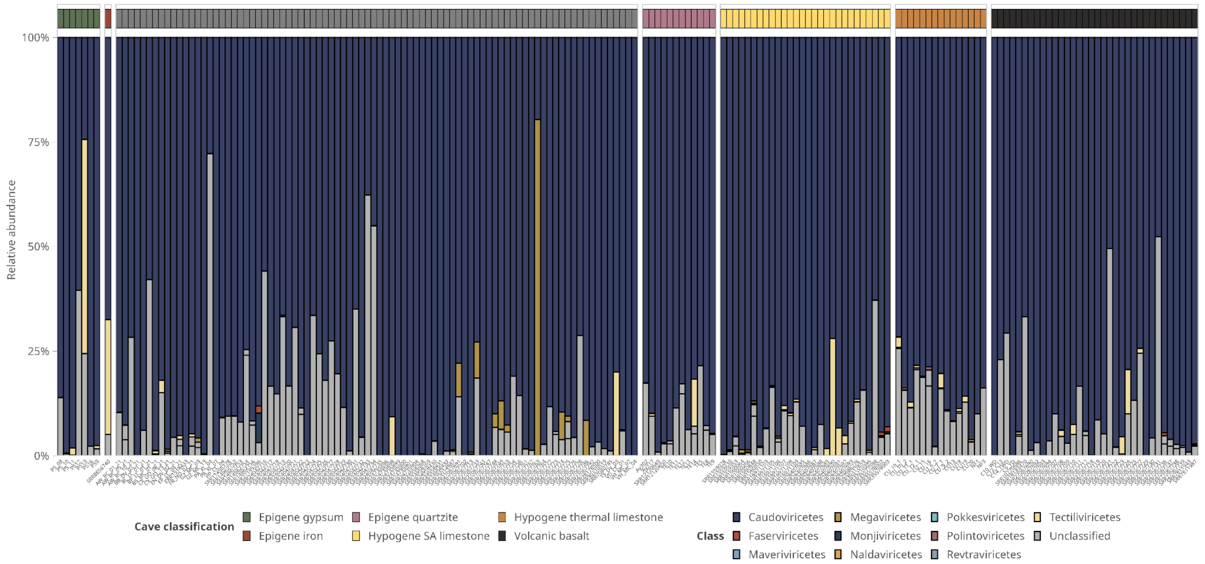


**Supplementary figure 5.** Diversity bar plot of taxonomic distribution of vOTUs across all samples

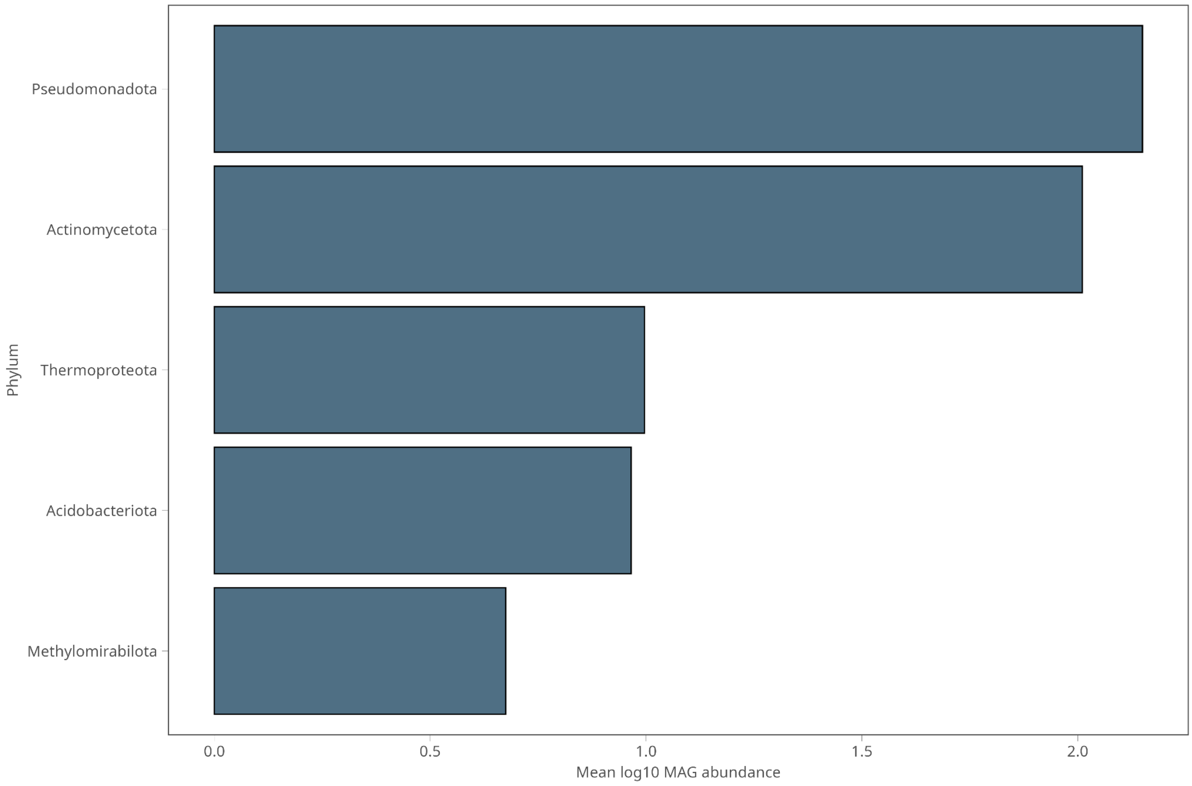


**Supplementary figure 6.** Mean log10 abundance of the microbial groups across all samples.

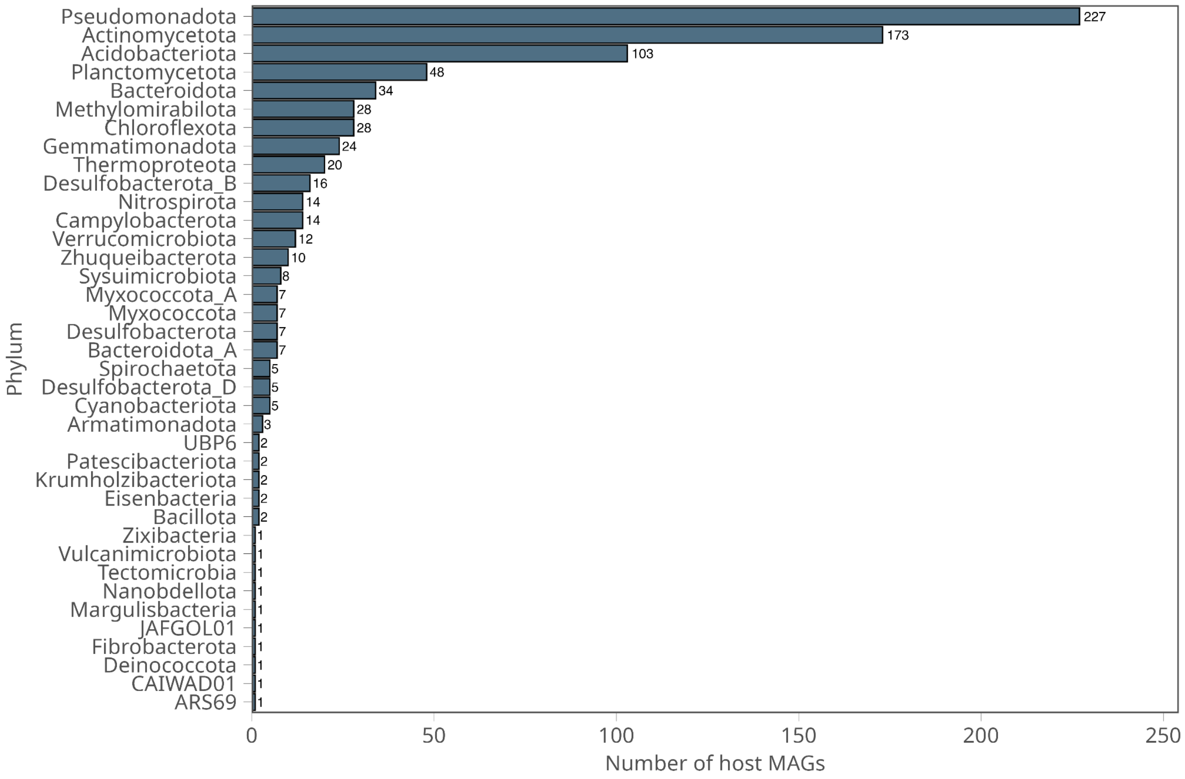


**Supplementary figure 7.** Nr of MAGs per bacterial phyla with viral links

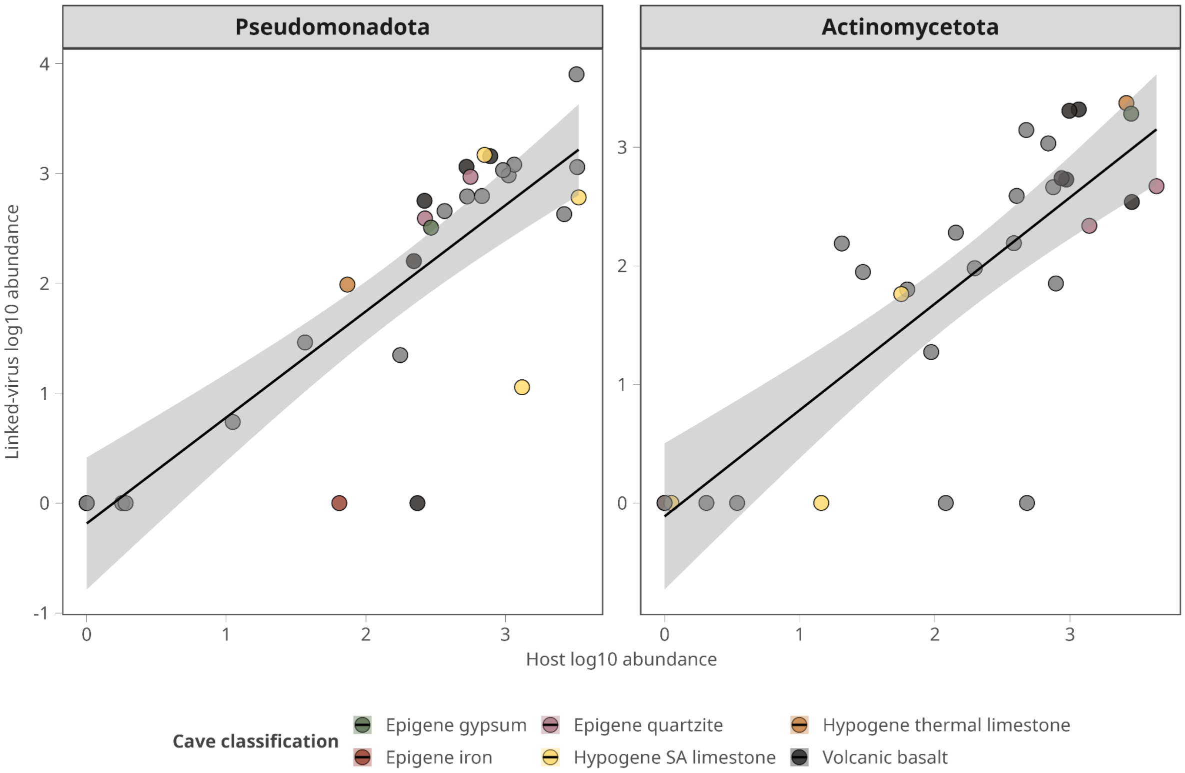


**Supplementary figure 8.** Correlation plot between the abundance of the vOTUs and their corresponding hosts belonging to the two most abundant phyla present in caves.

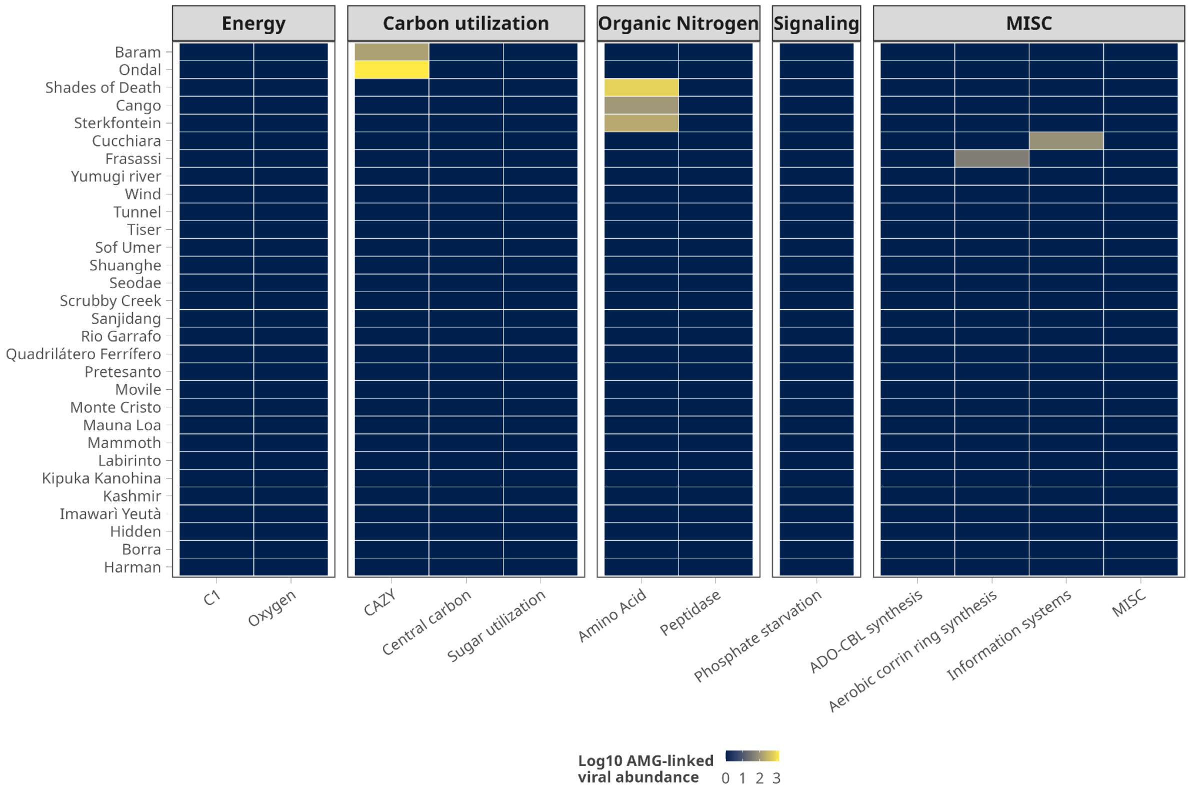


**Supplementary figure 9.** Heatmap depicting the abundance of the vOTUs annotated as having a lysogenic lifestyle and their corresponding auxiliary metabolic gene categories.
